# RAB3 phosphorylation by pathogenic LRRK2 impairs trafficking of synaptic vesicle precursors

**DOI:** 10.1101/2023.07.25.550521

**Authors:** Dan Dou, Jayne Aiken, Erika L.F. Holzbaur

**Affiliations:** Department of Physiology, Perelman School of Medicine, University of Pennsylvania, Philadelphia, PA 19104, USA; Aligning Science Across Parkinson’s (ASAP) Collaborative Research Network, Chevy Chase, MD 20815, USA; Neuroscience Graduate Group, University of Pennsylvania Perelman School of Medicine, Philadelphia, PA 19104, USA

## Abstract

Gain-of-function mutations in the *LRRK2* gene cause Parkinson’s disease (PD), characterized by debilitating motor and non-motor symptoms. Increased phosphorylation of a subset of RAB GTPases by LRRK2 is implicated in PD pathogenesis. We find that increased phosphorylation of RAB3A, a cardinal synaptic vesicle precursor (SVP) protein, disrupts anterograde axonal transport of SVPs in iPSC-derived human neurons (iNeurons) expressing hyperactive *LRRK2*-p.R1441H. Knockout of the opposing protein phosphatase 1H (*PPM1H*) in iNeurons phenocopies this effect. In these models, the compartmental distribution of synaptic proteins is altered; synaptophysin and synaptobrevin-2 become sequestered in the neuronal soma with decreased delivery to presynaptic sites along the axon. We find that RAB3A phosphorylation disrupts binding to the motor adapter MADD, potentially preventing formation of the RAB3A-MADD-KIF1A/1Bβ complex driving anterograde SVP transport. RAB3A hyperphosphorylation also disrupts interactions with RAB3GAP and RAB-GDI1. Our results reveal a mechanism by which pathogenic hyperactive LRRK2 may contribute to the altered synaptic homeostasis associated with characteristic non-motor and cognitive manifestations of PD.

**SUMMARY:** Dou et al. demonstrate that Parkinson’s disease-associated hyperactive LRRK2 decreases trafficking of synaptic vesicle proteins within neurons by disrupting regulation of the synaptic vesicle precursor protein RAB3A. Impaired delivery of synaptic proteins to presynaptic sites could contribute to progression of debilitating non-motor PD symptoms.

## INTRODUCTION

Parkinson’s disease (PD) is a devastating neurodegenerative disease that causes cardinal motor symptoms: rest tremor, rigidity, bradykinesia, and postural instability^1^. These are manifestations of the loss of select neuronal populations, most prominently dopaminergic neurons in the substantia nigra pars compacta (SNc). In addition to these motor symptoms, PD is clinically characterized by debilitating non-motor symptoms such as cognitive decline, dementia, sleep disturbance, and depression^1^, suggesting that pathogenic mechanisms may also alter synaptic transmission in a broader set of neuronal populations.

Autosomal dominant missense mutations in the leucine-rich repeat kinase 2 (*LRRK2*) gene are the most common genetic cause of PD, accounting for ∼5% of familial cases^2^. Furthermore, genome-wide association studies have implicated *LRRK2* noncoding variants in sporadic PD. Seven gain-of-function pathogenic mutations in *LRRK2* increase LRRK2 kinase activity, resulting in elevated phosphorylation of a subset of RAB GTPases (RABs)^3^. RABs coordinate vesicle trafficking by selectively associating with membrane compartments and recruiting effector proteins^4^. Mounting evidence shows that LRRK2-mediated phosphorylation of RABs alters their binding properties, either by introducing a new set of binding partners^5–9^ or by impairing interaction with previous partners^3, 10, 11^. Therefore, the relative activity of LRRK2 and its opposing protein phosphatase 1H (PPM1H) regulates RAB binding to effectors^12, 13^.

In recent work, we demonstrated that LRRK2-mediated RAB hyperphosphorylation has consequences for retrograde axonal transport of autophagic vesicles (AVs), disrupting an interplay of motor regulators in a manner scaling with magnitude of LRRK2 kinase activity^12^. Neurons require the directed transport of a wide range of distinct axonal cargoes to maintain homeostasis and synaptic function, in addition to AVs. Given the cognitive impairments and other non-motor manifestations of PD, an axonal cargo of particular interest is the synaptic vesicle precursor (SVP). SVPs arise in the neuronal soma and are transported anterogradely by kinesin-3 family members KIF1A and KIF1Bβ, carrying proteins that are fated for mature synaptic vesicles (SVs) at presynaptic sites^14–20^, including numerous *en passant* synapses populating the complex axonal arbor. These SV proteins are only recycled for a limited time before being targeted for degradation, necessitating robust delivery of new SVPs to replenish SV machinery^21, 22^.

Anterograde SVP transport is initiated by formation of a complex between RAB3, KIF1A/1Bβ, and a protein called “differentially expressed in normal and neoplastic cells/MAP kinase activating death domain” (DENN/MADD, henceforth referred to as MADD)^16, 18^. All three components of this complex are essential for the rapid, long-range transport of SVPs. MADD is a large ∼183 kDa protein that is also known as RAB3-GEP (guanine nucleotide exchange protein) due to its role as a GDP-GTP exchange factor (GEF) for RAB3^23^. Indeed, previous work has shown that the anterograde transport of SVPs depends on the GTP-bound state of RAB3^16, 24^.

RAB3 exists in four isoforms (RAB3A/B/C/D), with RAB3A being the most abundant in the brain cortex, although all four isoforms act redundantly in neurons^25, 26^. Importantly, all four isoforms are endogenous substrates of LRRK2^3, 10^ and are dephosphorylated by PPM1H^13^. However, the downstream consequences of LRRK2-mediated RAB3 phosphorylation on axonal transport have not been explored.

Here, we demonstrate that the hyperactive *LRRK2*-p.R1441H mutation reduces anterograde axonal flux of SVPs in gene-edited iPSC-derived human neurons (iNeurons). As an orthogonal model of RAB3 hyperphosphorylation, we show that knock-out (KO) of *PPM1H* phenocopies the effect of p.R1441H KI, indicating an important balance between LRRK2 and its opposing phosphatase for regulation of SVP transport. This transport deficit alters the distribution of SVP-associated proteins within the neuron, causing somal sequestration and decreased delivery to presynaptic sites in a heterologous synaptic model. Increasing levels of active RAB3 induced by overexpression of a GTP-locked mutant rescued the SVP transport deficit in p.R1441H KI iNeurons. Using a co-immunoprecipitation approach, we show that phosphorylation of RAB3A at the threonine 86 (T86) residue disrupts its binding to the motor adaptor protein MADD and the regulatory proteins RAB-GDI1 and RAB3GAP, but does not indiscriminately disrupt interactions with other known RAB3 binding partners. Together, our results uncover a mechanism by which pathogenic hyperactive LRRK2 mutations may contribute to synaptic dysfunction manifesting as debilitating motor and non-motor PD symptoms.

## RESULTS

### Endogenous LRRK2-p.R1441H impairs anterograde axonal transport of synaptic vesicle precursors

Point mutations at the p.R1441 hotspot (p.R1441C/G/H) in *LRRK2* are pathogenic for Parkinson’s disease (PD) and have high penetrance^27–30^. Previous reports have shown that these gain-of-function mutations induce hyperactivity of the LRRK2 kinase domain in multiple systems^3, 12, 31^ including neurogenin-2 (NGN2)-induced human neurons which express endogenous *LRRK2*^8, 32, 33^. One RAB GTPase that is phosphorylated by LRRK2 is RAB3A, which is associated with synaptic vesicle precursors (SVPs)^25^ and is essential for the anterograde axonal transport of SVPs^16, 18^. Given reports of altered binding properties of pRABs^3, 5–10^, we hypothesized that *LRRK2*-p.R1441H impairs anterograde axonal SVP transport via aberrant phosphorylation of RAB3A.

To test this hypothesis, we employed human induced pluripotent stem cells (iPSCs) with heterozygous KI of *LRRK2*-p.R1441H. These iPSCs had been gene-edited by the iPSC Neurodegenerative Disease Initiative (iNDI) at the NIH^34^ to introduce the p.R1441H mutation at the endogenous *LRRK2* locus of the KOLF2.1J parental line. Using tetracycline-inducible expression of *NGN2*, we differentiated these iPSCs into excitatory glutamatergic neurons (iNeurons)^35^. The resulting p.R1441H KI iNeurons exhibit elevated RAB phosphorylation, as previously shown using an antibody pan-specific to multiple phosphorylated RABs including RAB3A^12^. To assess SVP trafficking in these mutant iNeurons and corresponding WT control iNeurons, we live-imaged SVPs labeled by the fluorescent reporter mScarlet-synaptophysin (SYP) (Figure 1A). We imaged each neuron at the proximal axon in order to limit variability caused by axonal branchpoints, at a distance approximately 100 μm from the soma in order to avoid the axonal initial segment. To more clearly visualize SYP+ vesicles entering the imaged axonal region, we photobleached the field of view prior to imaging to deplete pre-existing mScarlet-SYP signal in the axon (Figure 1A).

**Figure 1.**
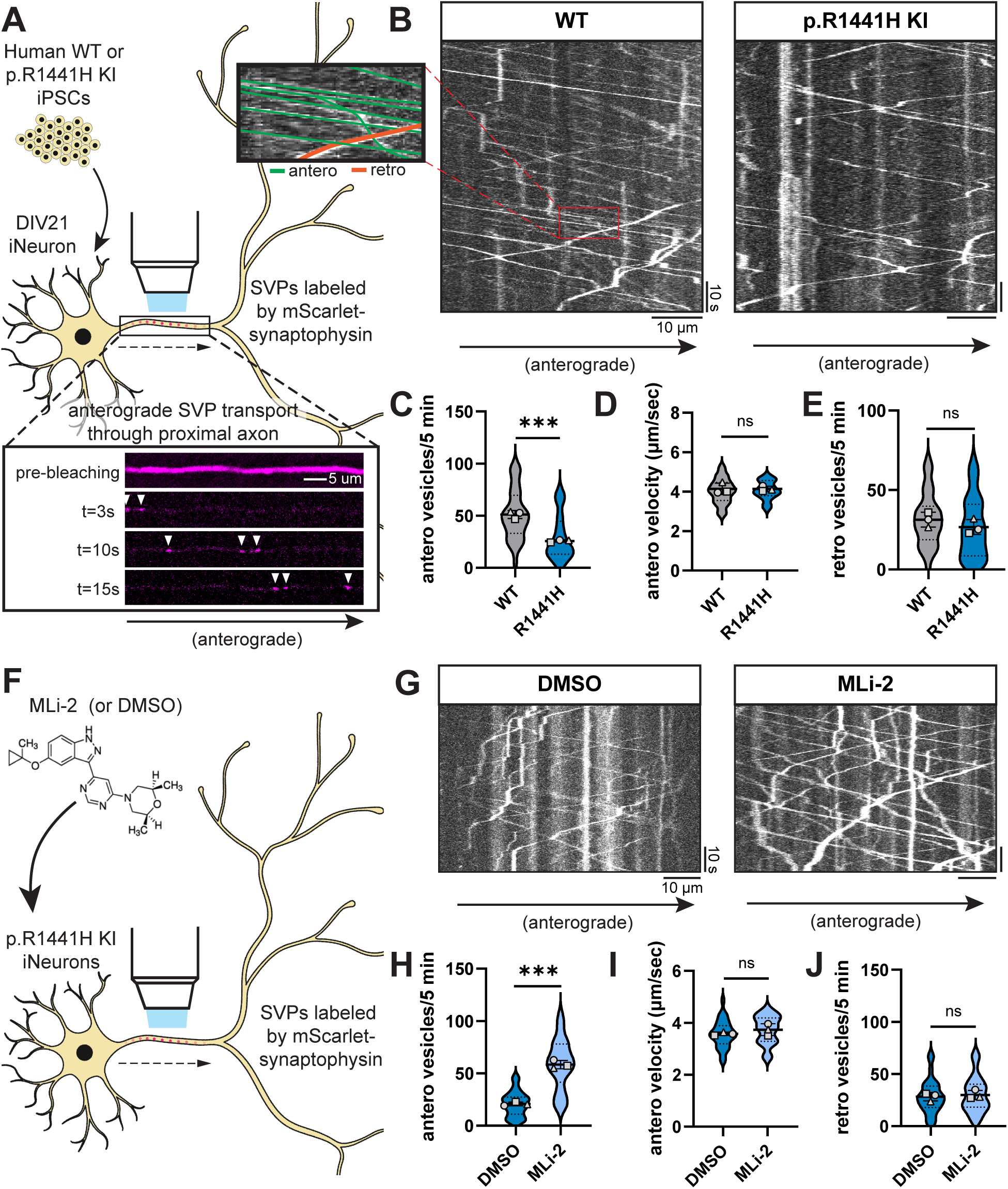
*LRRK2*-p.R1441H knockin causes kinase-dependent decrease in anterograde axonal SVP flux. (A) Inset, below: example time lapse images of mScarlet-SYP+ vesicles in the proximal axon of a DIV21 WT iPSC-derived neuron (iNeuron). Cytoplasmic mScarlet-SYP signal was photobleached at t=0s. (B) Kymographs of axonal mScarlet-SYP+ vesicles in DIV21 WT and p.R1441H KI iNeurons. Inset, left: example traces of anterograde and retrograde SYP+ vesicles. (C-E) Anterograde flux (C), anterograde velocity (D), and retrograde flux (E) of SYP+ vesicles in WT and p.R1441H KI iNeurons (n = 24-33 neurons from 3 independent experiments; ns, not significant, p=0.8628 for antero velocity, p=0.2978 for retro vesicles; ***p<0.001; linear mixed effects model). (F) Cartoon depicting p.R1441H KI iNeuron treated overnight with DMSO or 100 nM MLi-2. (G) Kymographs of axonal mScarlet-SYP+ vesicles in p.R1441H KI iNeurons treated with DMSO or MLi-2. (H-J) Anterograde flux (H), anterograde velocity (I), and retrograde flux (J) of SYP+ vesicles in p.R1441H KI iNeurons treated with DMSO or MLi-2 (n = 29-30 neurons from 3 independent experiments; ns, not significant, p=0.2344 for antero velocity, p=0.6735 for retro vesicles; ***p<0.001; linear mixed effects model). Scatter plot points indicate the means of three independent experiments, and error bars show mean ± SD of these points.

In WT iNeurons we observed rapid, highly-processive SVPs traveling in the anterograde direction (Figure 1B). To accommodate the high speed of these vesicles, we imaged each axon at 5 frames per second for 5 minutes. We observed the anterograde population of SYP+ vesicles to be more numerous, rapid, and processive than the retrograde population (Figure 1B, inset), consistent with our previous observations in WT primary mouse hippocampal neurons^22^ and in WT iNeurons from a different parental line^36^ (Aiken and Holzbaur, 2023; manuscript in preparation). In *LRRK2*-p.R1441H KI iNeurons, we observed a significant decrease in anterograde SVP flux (Figure 1B,C). However, there was no effect on the velocity of anterograde vesicles (Figure 1D), indicating that SVPs that entered the axon were transported normally. We did not observe a change in the flux of retrograde SYP+ vesicles (Figure 1E), suggesting that expression of mutant *LRRK2* specifically affected the anterograde population.

To confirm that this effect is dependent on LRRK2 kinase activity, we applied the selective LRRK2 kinase inhibitor MLi-2^37^ to p.R1441H KI iNeurons (Figure 1F). Overnight treatment with 100 nM MLi-2 rescued anterograde SVP flux (Figure 1G,H) without affecting anterograde velocity (Figure 1I). MLi-2 treatment had no effect on flux of retrograde SYP+ vesicles (Figure 1J). In parallel experiments, we also examined SVP flux in i^3^Neurons^8, 38, 39^ gene-edited from the WTC11 parental line to express the common pathological p.G2019S variant of *LRRK2*. This mutation also hyperactivates kinase activity but to a lesser extent than the p.R1441H mutation^3, 12, 31^. In p.G2019S KI i^3^Neurons, we found that overnight treatment with 100 nM MLi-2 increased anterograde SVP flux compared to DMSO treatment (Figure S1A-C), albeit to a lesser extent than was observed in iNeurons expressing p.R1441H.

Together, these results show a kinase activity-dependent decrease in anterograde axonal SVP flux caused by hyperactive LRRK2. Importantly, we did not observe altered anterograde SVP velocity, nor did we observe altered retrograde flux, suggesting that p.R1441H’s effect is specific to decreasing the number of SVPs that enter the axon from the soma.

### *PPM1H* KO phenocopies the effect of hyperactive LRRK2 on anterograde SVP flux in the axon

PPM1H is a phosphatase that opposes the activity of LRRK2 through dephosphorylation of RAB GTPases (Figure 2A)^13, 40, 41^. Therefore, the balance of LRRK2 and PPM1H activity has the potential to regulate neuronal pathways, including transport of axonal cargoes, by modulating RAB phosphorylation levels^12^. We previously generated *PPM1H* KO iPSCs from the same KOLF2.1J parental line as the p.R1441H KI iPSCs used here^12^. To determine whether loss of PPM1H would phenocopy the effect of hyperactive *LRRK2*-p.R1441H on anterograde SVP flux, we compared mScarlet-SYP motility in WT and *PPM1H* KO iNeurons (Figure 2B). Indeed, we observed decreased anterograde axonal flux of the SYP+ population upon loss of PPM1H (Figure 2C,D). Similar to observations in p.R1441H KI neurons, there was no change in anterograde velocity in *PPM1H* KO cells (Figure 2E). While we did observe a significant decrease in retrograde SYP+ vesicles in *PPM1H* KO iNeurons (Figure 2F), the size of the effect was much smaller than the effect on the anterograde population (Figure 2D).

**Figure 2.**
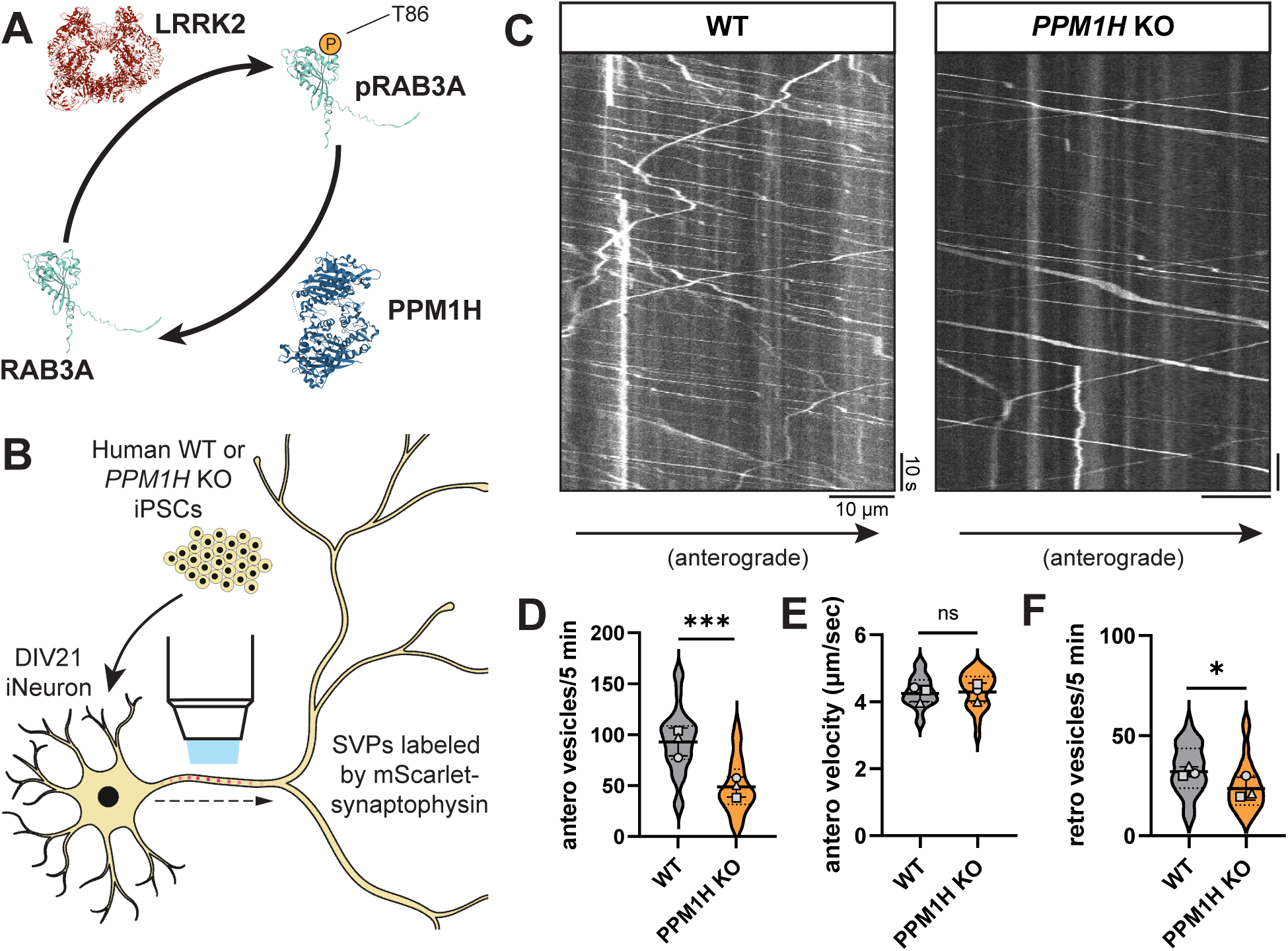
Loss of LRRK2-opposing PPM1H decreases anterograde axonal SVP flux. (A) Schematic depicting opposing regulation of RAB3A phosphorylation state at threonine 86 by LRRK2 kinase and PPM1H phosphatase. Protein structures: LRRK2 (PDB: 7LHT), PPM1H (PDB: 7L4J), RAB3A (AlphaFold prediction^40, 41^). (B) Cartoon depicting WT or *PPM1H* KO iNeuron expressing mScarlet-SYP. (C) Kymographs of axonal mScarlet-SYP+ vesicles in DIV21 WT and *PPM1H* KO iNeurons. (D-F) Anterograde flux (D), anterograde velocity (E), and retrograde flux (F) of SYP+ vesicles in WT and *PPM1H* KO iNeurons (n = 28 neurons from 3 independent experiments; ns, not significant, p>0.6872; *p=0.0119; ***p<0.001; linear mixed effects model). Scatter plot points indicate the means of three independent experiments, and error bars show mean ± SD of these points.

In sum, these results from orthogonal models indicate that either hyperactive LRRK2 activity (Figure 1) or knockout of the opposing phosphatase (Figure 2) leads to decreased anterograde flux of SVPs.

### RAB hyperphosphorylation alters compartmental distribution of SVP-associated proteins

SVP transport has the important role of replenishing presynaptic sites with synaptic vesicle (SV) proteins^21, 22^. We next asked whether decreased anterograde transport of SVPs caused by RAB hyperphosphorylation has consequences on distribution of synaptic proteins within the neuron. Specifically, we interrogated the localization of two SV proteins known to be trafficked with SVPs and delivered to presynaptic sites: SYP and synaptobrevin-2 (SYB2).

To examine somal content, we stained endogenous SYP and SYB2 in p.R1441H KI and *PPM1H* KO iNeurons, as well as control WT KOLF2.1J iNeurons. An antibody to endogenous microtubule-associated protein-2 (MAP2) signal was used to visualize the somatodendritic compartment (Figure 3A). In both RAB-hyperphosphorylated conditions, we detected significant increases in the somal intensity of SYP (Figure 3A,B) and SYB2 (Figure 3A,C), with no associated changes in somal size as quantified by the total area of somal MAP2 signal (Figure S2A). Western blots of neuronal lysates showed no evidence of changed levels of SYP, SYB2, or RAB3A (Figure S2B-D).

**Figure 3.**
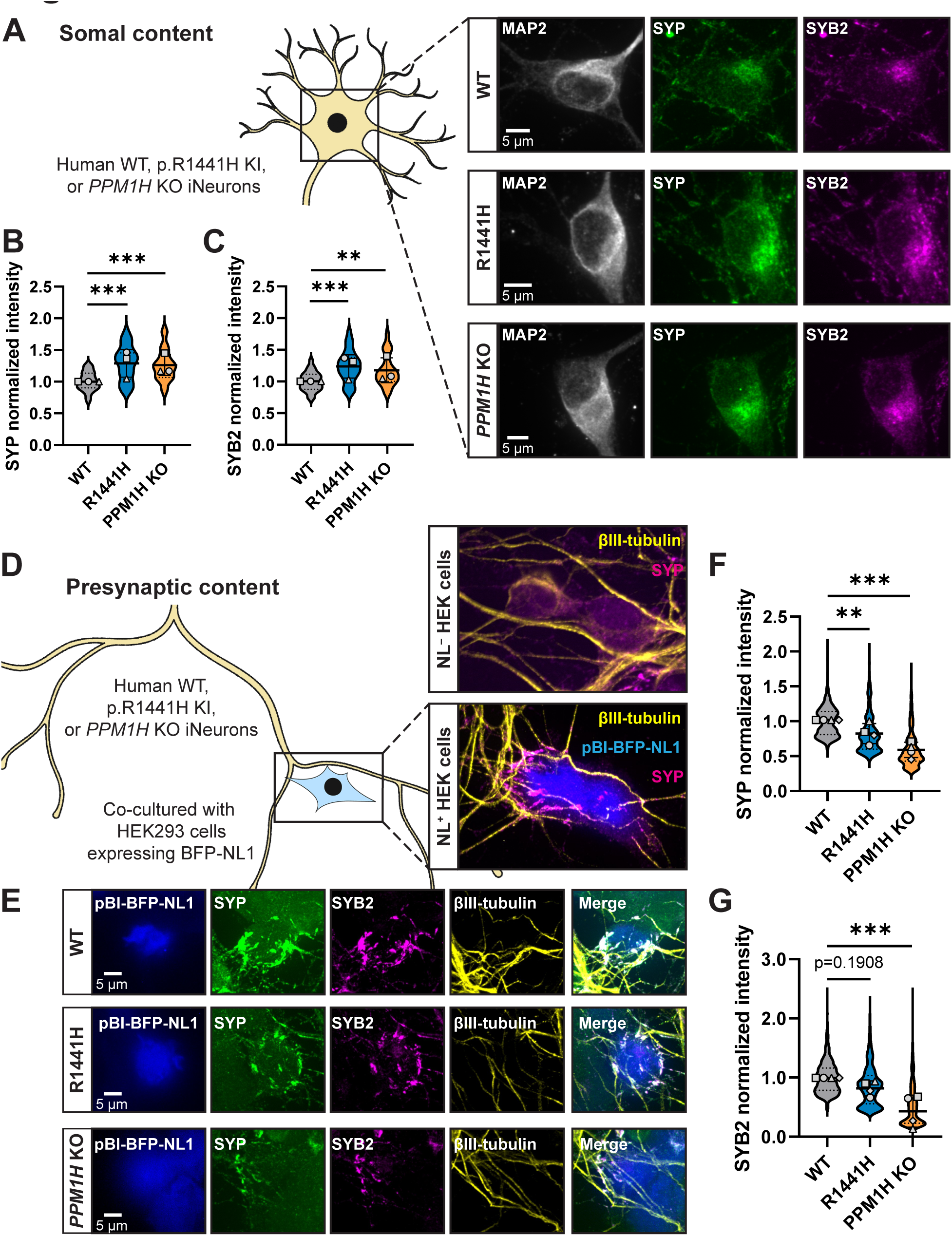
Delivery of synaptic vesicle proteins from soma to presynaptic sites is disrupted by RAB hyperphosphorylation. (A) Representative images of DIV14 WT, p.R1441H KI, and *PPM1H* KO iNeuron somas, stained for endogenous MAP2, SYP, and SYB2. (B,C) Normalized somal intensity (mean grey value) of SYP (B) and SYB2 (C) in WT, p.R1441H KI, and *PPM1H* KO iNeurons (n = 24 neurons from 3 independent experiments; **p=0.0027; ***p<0.001; linear mixed effects model). (D) Cartoon depicting heterologous synapse model set-up, where HEK293 cells expressing pBI-BFP-NL1 were introduced in co-culture to WT, p.R1441H KI, or *PPM1H* KO iNeurons. Dishes were imaged at iNeuron DIV14, at which point βIII-tubulin+ iNeuron axons had selectively formed presynapses with NL1+ transfected HEK cells (inset, bottom), but not untransfected HEK cells (inset, top). (E) Representative images of heterologously modeled synapses including HEK293 cells expressing pBI-BFP-NL and WT, p.R1441H KI, or *PPM1H* KO iNeurons, stained for endogenous SYP, SYB2, and βIII-tubulin. (F) Normalized intensity of SYP at heterologous presynaptic sites in WT, p.R1441H KI, and *PPM1H* KO iNeurons (n = 492-840 presynaptic puncta in 17-20 fields of view from 4 independent experiments; **p=0.0086; ***p<0.001; linear mixed effects model). (G) Normalized intensity of SYB2 at at heterologous presynaptic sites in WT, p.R1441H KI, and *PPM1H* KO iNeurons (mean ± SD; n = 1018-1512 presynaptic puncta in 16-20 fields of view from 4 independent experiments; ***p<0.001; linear mixed effects model). Scatter plot points indicate the means of 3-4 independent experiments, and error bars show mean ± SD of these points.

Proteins fated for SVPs are believed to be sorted at the trans-Golgi network (TGN) ^42–44^. We next explored whether a portion of somal SYP may be sequestered at the TGN in the context of elevated RAB3A phosphorylation. WT, p.R1441H KI, and *PPM1H* KO iNeurons were stained for endogenous SYP and golgin-97, a TGN marker (Figure S2E). Again, MAP2 signal was used to visualize the somatodendritric compartment. In all three conditions, we observed that SYP intensity is enriched at the TGN relative to whole soma (Figure S2F). Compared to WT neurons, *PPM1H* KO neurons displayed increased SYP intensity that co-localized with golgin-97 (Figure S2F). However, we did not detect this effect in p.R1441H KI iNeurons (Figure S2F).

Next, we sought to determine whether hyperphosphorylation of RABs disrupts delivery of synaptic proteins to presynaptic sites. To accomplish this, we employed a recently developed heterologous synapse model for human neurons that allows unambiguous analysis of trafficking to the presynaptic compartment^36^ (Aiken and Holzbaur, 2023; manuscript in preparation), introducing non-neuronal human embryonic kidney (HEK) 293 cells expressing the postsynaptic ligand neuroligin-1 (*NL1*) into co-culture with iNeurons (Figure 3D). Within 24 hours of introducing HEK cells, iNeuron axons specifically recognize *NL1*-expressing HEK cells and form connections where presynaptic proteins accumulate (Figure 3D, inset). This system provides both spatial and temporal control for quantification of SVP-associated protein accumulation. These heterologous presynapses contain SYP, SYB2, synapsin I/II (SYN), VGLUT1, and SVs that cycle upon neuronal depolarization^36^ (Aiken and Holzbaur, 2023; manuscript in preparation). In heterologous synaptic cultures stained for endogenous SYP and SYB2 (Figure 3E), quantification revealed significantly decreased accumulation of both SYP (Figure 3F) and SYB2 (Figure 3G) at presynapses in *PPM1H* KO iNeurons relative to WT. In p.R1441H KI iNeurons, SYP presynaptic accumulation was also decreased, with SYB2 presynaptic accumulation trending downward but not achieving statistical significance (Figure 3F-G).

Together, these experiments show that the compartmental distribution of two different SVP-associated proteins is disrupted in p.R1441H KI and *PPM1H* KO iNeurons, with increased somal abundance and decreased presynaptic content. Together with our live-imaging findings, these data are consistent with somal sequestration and impaired presynaptic delivery of SVP-associated proteins as a consequence of disrupted anterograde SVP transport.

### Phosphorylation of RAB3A impairs interaction with motor adaptor protein MADD

MADD has been previously shown to be essential for transport of RAB3-containing SVPs by KIF1A and KIF1Bβ^16^. The same study showed that MADD directly interacts with KIF1A/1Bβ and RAB3. A more recent study further demonstrated that MADD interacts selectively with SVPs, not other axonal cargoes including dense core vesicles (DCVs) and lysosomes^18^. MADD is also known as RAB3-GEP (guanine nucleotide exchange protein) due to its role as a GDP-GTP exchange factor (GEF) for RAB3^23^. Notably, anterograde SVP transport has been shown to depend on the GTP-bound state of RAB3^16, 24^. To probe the mechanism underlying *LRRK2-*p.R1441H’s effect on anterograde axonal SVP flux, we first tested whether overexpressing the GTP-locked mutant of *RAB3* would rescue the deficit. Of the four RAB3 isoforms, we chose to focus on the best-characterized isoform, RAB3A, which is also the most abundant in the cortex^25, 26^. We found that transient expression of the glutamine-to-leucine (Q81L) mutant RAB3A, predicted to lock RAB3A into a GTP-bound state (Figure 4A)^40, 41^, abolished the inhibitory effect of the hyperactive LRRK2 mutant p.R1441H KI on anterograde SVP flux (Figures 1B-C,4B-D).

**Figure 4.**
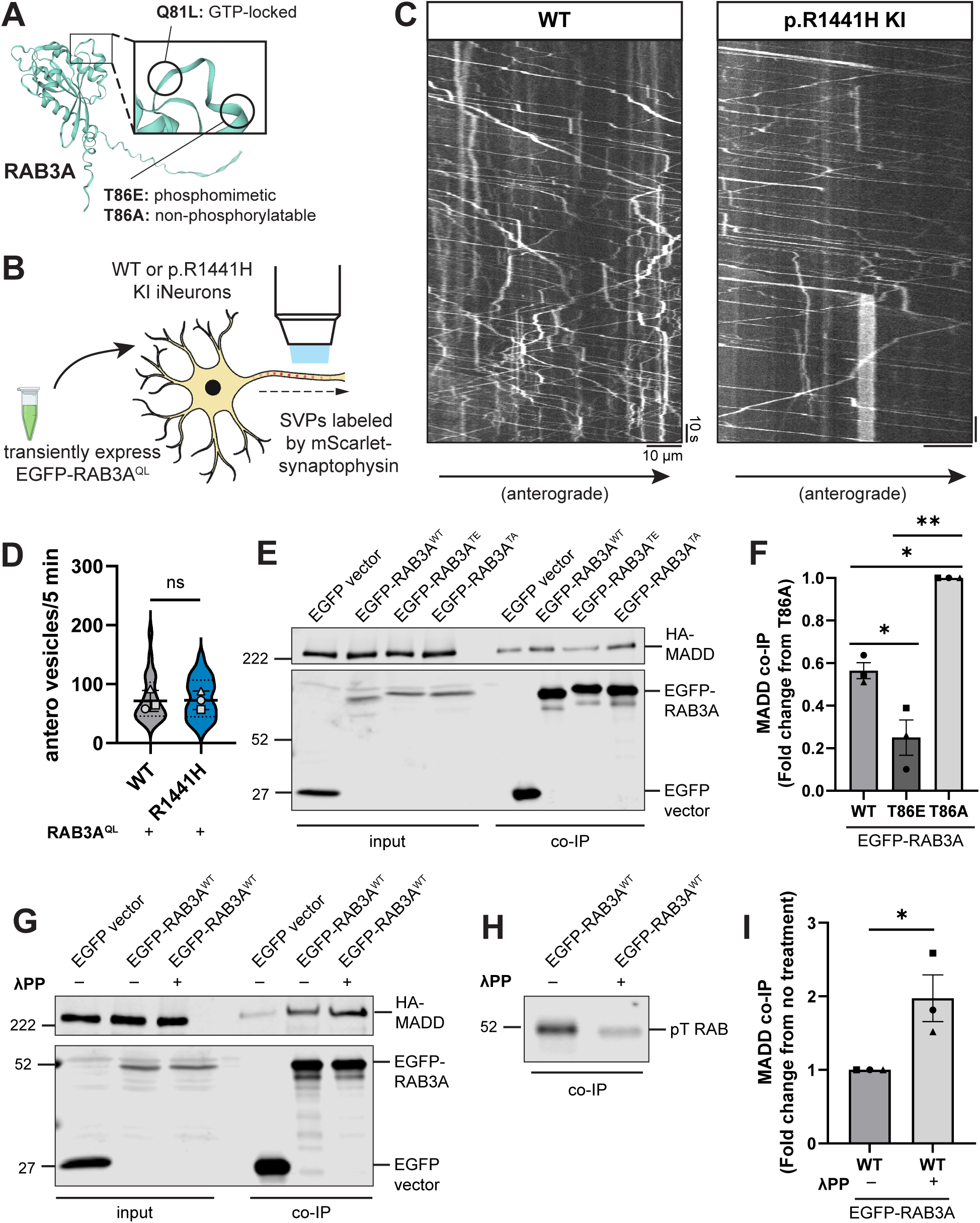
Phosphorylation of RAB3A disrupts binding to motor adapter MADD. (A) AlphaFold prediction^40, 41^ of RAB3A. Inset illustrates locations of site-directed mutagenesis for RAB3A mutants used in this study. (B) Cartoon depicting experimental approach. EGFP-RAB3A^QL^ was transiently expressed in DIV21 WT or p.R1441H KI iNeurons expressing mScarlet-SYP. (C) Kymographs of axonal mScarlet-SYP+ vesicles in DIV21 WT and p.R1441H KI iNeurons, transiently expressing EGFP-RAB3A^QL^. (D) Anterograde flux of SYP+ vesicles in WT and p.R1441H KI iNeurons, transiently expressing EGFP-RAB3A^QL^ (n = 23-24 neurons from 3 independent experiments; ns, not significant, p=0.8993 linear mixed effects model). Scatter plot points indicate the means of 3 independent experiments, and error bars show mean ± SD of these points. (E,F) Example immunoblot and quantification of MADD co-immunoprecipitation by RAB3A^WT^, RAB3A^TE^, or RAB3A^TA^, co-expressed in HEK293T cells (mean ± SEM; n = 3 independent experiments; *p=0.0405 for WT vs T86E, p=0.0140 for WT vs T86A; **p=0.0018; one-way ANOVA with Tukey’s multiple comparisons test). (G) Example immunoblot of MADD co-immunoprecipitation by RAB3A^WT^, co-expressed in HEK293T cells, with or without 30 minutes treatment of lysate with lambda protein phosphatase (λPP; 200 units λPP per 50 μL reaction volume). (H) Example immunoblot of phosphothreonine RAB in bound fraction of MADD co-immunoprecipitation by RAB3A^WT^, with or without λPP treatment of lysate. (I) Quantification of MADD co-immunoprecipitation by RAB3A^WT^, co-expressed in HEK293T cells, with or without 30 minutes treatment of lysate with λPP (mean ± SEM; n = 3 independent experiments; *p=0.0303; unpaired t test). For all co-IP experiments shown, samples were processed and immunoblotted in parallel.

It has previously been shown that LRRK2-mediated phosphorylation of RAB8A disrupts its ability to bind to RABIN8, its cognate GEF^3, 11^. We therefore investigated whether the known interaction between RAB3A and MADD is altered by RAB3A phosphorylation. To test this, we co-expressed HA-MADD in HEK293 cells with EGFP-labeled RAB3A, with or without point mutations at the threonine residue that is phosphorylated by LRRK2 (Figure 4A)^40, 41^. Consistent with previous work^16^, RAB3A^WT^ co-immunoprecipitated with MADD (Figure 4E). Threonine-to-alanine (T86A) mutant RAB3A, predicted to be non-phosphorylatable, exhibited the highest binding affinity to MADD (Figure 4E,F). Threonine-to-glutamic acid (T86E) mutant RAB3A, predicted to be a phosphomimetic, bound more weakly to MADD than RAB3A^WT^ (Figure 4E,F).

Compared to the T86A and T86E mutants, RAB3A^WT^ exhibited an intermediate binding affinity with MADD (Figure 4E,F). This raised the intriguing possibility that a fraction of the transiently-expressed EGFP-RAB3A^WT^ was phosphorylated by HEK cell endogenous LRRK2^WT^ and thus exhibited an impaired ability to bind MADD. To confirm that phosphorylation of RAB3A^WT^ affects MADD binding, we applied lambda protein phosphatase (λPP) to lysates prior to co-immunoprecipitation of EGFP-RAB3A^WT^ and HA-MADD (Figure 4G). λPP treatment effectively decreased levels of phosphorylated EGFP-RAB3A^WT^ in the bound fraction (Figure 4H) and increased binding to HA-MADD (Figure 4G,I).

In sum, our results show that interaction between RAB3A and MADD is impaired by RAB3A phosphorylation at the T86 residue. Given MADD’s dual role as a GEF for RAB3A and a motor adaptor for RAB3A-positive SVPs, these data suggest that impaired RAB3A-MADD interaction contributes to the deficit of anterograde SVP flux in p.R1441H KI iNeurons (Figure 1A-C), which can be rescued by overexpression of GTP-locked RAB3A (Figure 4D).

### Phosphorylation of RAB3A impairs interactions with RAB-GDI1 and RAB3GAP but not RIM2 or synapsin

Multiple regulatory proteins determine RAB GTPase localization and binding state^4^. GEFs such as MADD recruit and drive conversion of RABs to the active GTP-bound state at membranes^23^. Each RAB functions at specific membranes, contributing to membrane identity by selective effector recruitment^45^. However, cytosolic GDP-bound RABs have been shown to be “promiscuous” in terms of their ability to enter membranes belonging to a wide range of intracellular organelles, where they may fail to encounter their cognate GEF^46^. Two regulatory proteins called RAB3 GTPase-activating protein (RAB3GAP) and RAB GDP dissociation inhibitor-1 (RAB-GDI1) act in concert to retrieve RABs from inappropriate membranes. RAB3GAP converts GTP-bound RABs to the GDP-bound state, and RAB-GDI1 serves as a chaperone to return GDP-bound RABs from membranes to the cytosol^4^. Dysregulation of RAB GTP binding state may therefore alter subcellular RAB localization and contribute to reduced effective availability of RABs^4, 47^.

Previous work showed that phosphomimetic mutant RAB GTPases fail to bind RAB-GDI1, and that this is also true for directly-phosphorylated RAB8A^WT^ and RAB12^WT^ ^3, 10, 48^. Consistent with these findings, in co-immunoprecipitation experiments we observed that the phosphomimetic T86E mutation abolished the interaction between EGFP-RAB3A and endogenous RAB-GDI1, compared to the non-phosphorylatable T86A mutant (Figure 5A,C). Similar to the RAB3A-MADD interaction (Figure 4E-F), RAB3A^WT^ exhibited intermediate binding affinity with GDI (Figure 5A,C). Notably, de-phosphorylation of RAB3A^WT^ by λPP treatment (Figure 5B) significantly increased binding to RAB-GDI1 (Figure 5A,C), confirming that direct RAB3A phosphorylation disrupts the interaction between RAB3A and RAB-GDI1.

**Figure 5.**
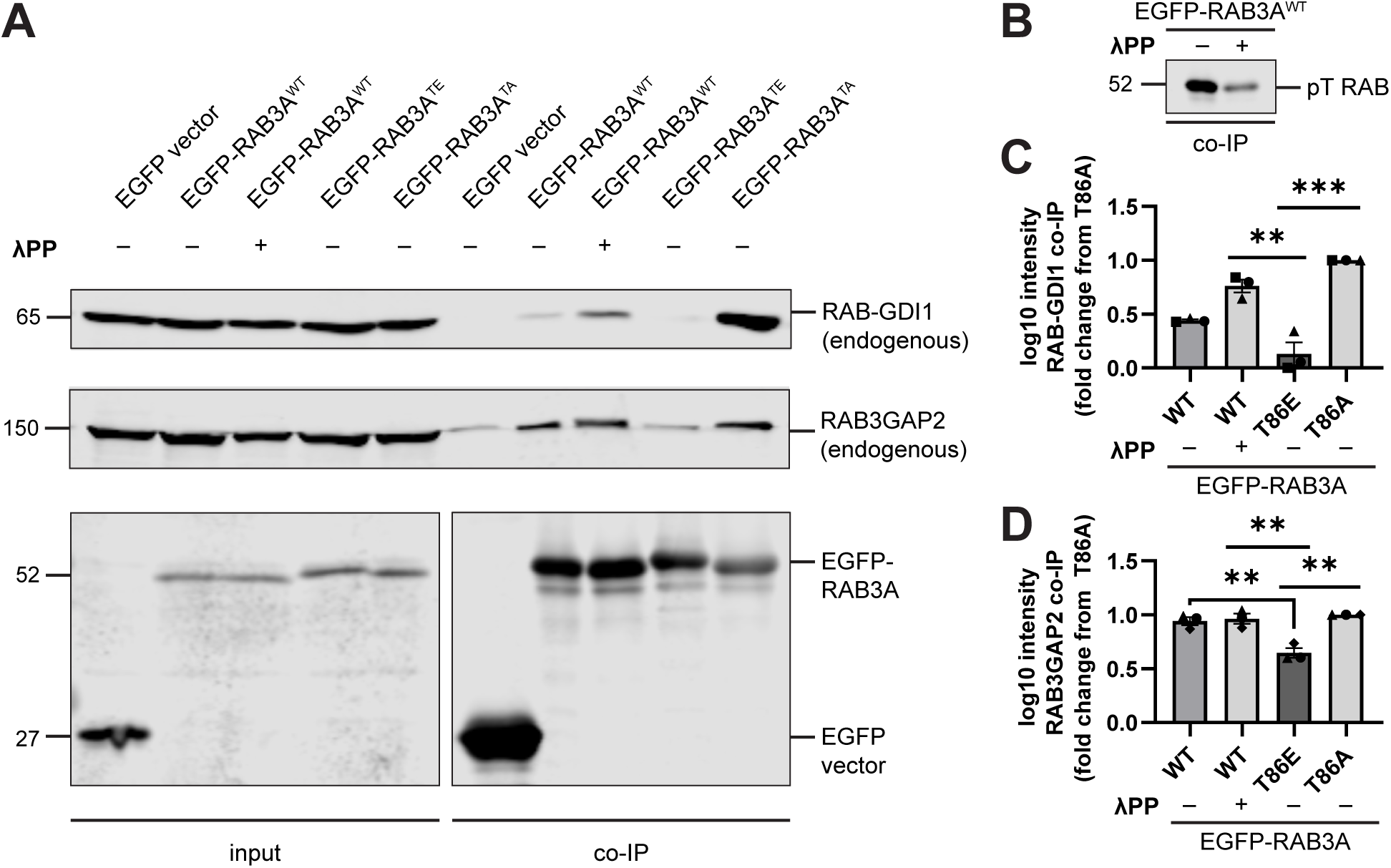
Phosphorylation of RAB3A disrupts binding to RAB cycle regulators RAB-GDI1 and RAB3GAP. (A) Example immunoblot of endogenous RAB3GAP2 and endogenous RAB-GDI1 co-immunoprecipitation by RAB3A^WT^, RAB3A^TE^, or RAB3A^TA^, expressed in HEK293T cells, with or without 30 minutes treatment of lysate with lambda protein phosphatase (λPP; 200 units λPP per 50 μL reaction volume). Lower panel is separated for alignment purposes; no lanes that included sample were excluded. (B) Example immunoblot of phosphothreonine RAB in bound fraction of endogenous RAB-GDI1 and RAB3GAP2 co-immunoprecipitation by RAB3A^WT^, with or without λPP treatment of lysate. (C) Quantification of endogenous RAB-GDI1 co-immunoprecipitation by RAB3A^WT^, RAB3A^TE^, or RAB3A^TA^, expressed in HEK293T cells, with or without 30 minutes treatment of lysate with λPP (mean ± SEM; n = 3 independent experiments; **p<0.0048; ***p<0.001; one-way ANOVA with Tukey’s multiple comparisons test). (D) Quantification of endogenous RAB3GAP2 co-immunoprecipitation by RAB3A^WT^, RAB3A^TE^, or RAB3A^TA^, expressed in HEK293T cells, with or without 30 minutes treatment of lysate with λPP (mean ± SEM; n = 3 independent experiments; **p<0.0094; one-way ANOVA with Tukey’s multiple comparisons test). For all co-IP experiments shown, samples were processed and immunoblotted in parallel.

In the same set of co-immunoprecipitation experiments, we also explored whether phosphorylation of RAB3A disrupts binding to RAB3GAP, quantified with an antibody for the non-catalytic subunit RAB3GAP2. We noted greater non-specific binding of RAB3GAP than GDI1 to the EGFP vector (Figure 5A), which was subtracted prior to quantification. We observed that binding of EGFP-RAB3A to RAB3GAP was decreased by the phosphomimetic T86E mutation compared to both RAB3A^WT^ and RAB3A^TA^ (Figure 5A,D). Interestingly, while the T86E mutation strongly disrupted the RAB3A-GDI1 interaction (Figure 5A,C), this mutation had a more moderate effect on the RAB3A-RAB3GAP interaction (Figure 5A,D).

RAB3A has been implicated in mechanisms of SV release, acting in concert with effector proteins^49–51^. Given the effect of RAB3A phosphorylation on binding to MADD, RAB-GDI1, and RAB3GAP, we wondered if RAB3A phosphorylation indiscriminately impaired interaction with all of its effectors. We therefore tested phosphomimetic mutant RAB3A binding to RAB3A-interacting molecule 2 (RIM2) and synapsin, two presynaptic proteins that have been shown to act as RAB3A effectors^50, 52^. In agreement with these previous reports, we observed pulldown of both RIM2 (Figure S3A) and synapsin (Figure S3B) by RAB3A^WT^. However, neither the phosphomimetic T86E mutation nor the nonphosphorylatable T86A mutation affected RAB3A binding to either RIM2 or synapsin, in contrast to MADD, GDI, and RAB3GAP.

In summary, our results show that RAB3A phosphorylation at T86 selectively regulates binding to a subset of partners. Binding to the regulatory chaperone protein RAB-GDI1 was strongly impaired, while moderate disruption was observed for binding to the motor adapter MADD and the regulatory protein RAB3GAP, and no effects were observed on binding to either RIM2 or synapsin.

## DISCUSSION

PD-linked pathogenic mutations in LRRK2 hyperphosphorylate a subset of RAB GTPases, with a growing body of evidence linking this posttranslational modification to altered interactions with effectors. Though the LRRK2 substrate RAB3A has long been known to be essential for axonal SVP transport, the consequences of RAB3A phosphorylation on this trafficking pathway have not previously been explored. Here, we show that in two iNeuron models of RAB hyperphosphorylation (*LRRK2*-p.R1441H KI and *PPM1H* KO), we observe impaired anterograde flux of SVPs (Figure 1,2). We find that RAB3A phosphorylation at T86 disrupts binding to the motor adapter protein MADD (Figure 4). Phosphorylation of RAB3A also alters binding affinity to the regulatory proteins RAB3GAP and RAB-GDI1 (Figure 5). Our findings support a model where pathogenic hyperactive LRRK2 causes dysregulated binding of RAB3A in the neuronal soma, including impairment of the formation of the RAB3A-MADD-KIF1A/Bβ complex (Figure 6). We hypothesize that this contributes to the reduced availability of RAB3 to stimulate anterograde SVP transport. Consistent with this hypothesis, we find that the compartmental distribution of SVP-associated proteins is disrupted within neurons with hyperphosphorylated RABs, manifesting as increased somal content and decreased delivery to presynaptic sites along the axon (Figure 3).

**Figure 6.**
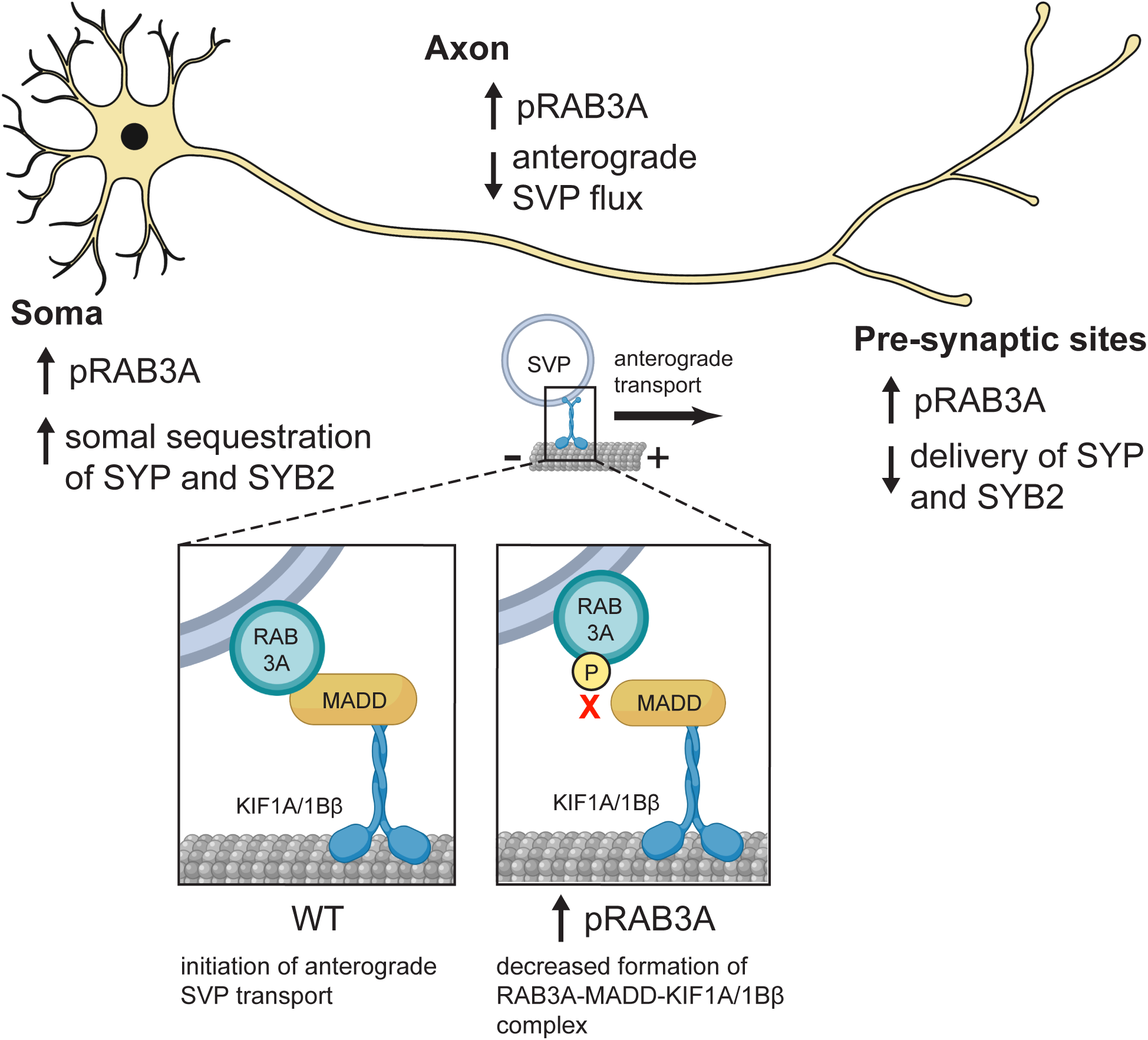
Model: Dysregulated pRAB3A binding disrupts axonal transport of SVPs and distribution of synaptic proteins. In the neuronal soma, increased phosphorylation of RAB3A by hyperactive LRRK2 results in impaired formation of the RAB3A-MADD-KIF1A/1Bβ motor complex (inset) that is necessary for anterograde transport of SVPs out of the soma. As a result, there is increased somal sequestration of SYP and SYB2, and decreased anterograde SVP flux in the axon. Consequently, decreased SYP and SYB2 is delivered to presynaptic sites.

Our previous work linked hyperactive LRRK2 mutations to the disruption of the retrograde axonal transport of AVs^8, 12^, most likely mediated by RAB10 and/or RAB35^5, 6, 13, 53^. In contrast, hyperactive LRRK2 does not alter the axonal transport of mitochondria^12^, consistent with our current understanding that there is no known role for RABs in regulating the axonal transport of these organelles. Given the altered transport of RAB3A+ SVPs, our findings indicate a high degree of RAB-dependent selectivity for which axonal cargoes are perturbed by pathogenic hyperactive LRRK2. Furthermore, because *PPM1H* KO phenocopies these transport defects, this implies that the balance between LRRK2^WT^ and PPM1H may regulate transport of both SVPs and autophagosomes under physiologic conditions. As at least ten different RABs are endogenously phosphorylated by LRRK2^10^, it remains to be explored whether other axonal cargoes rely on RAB-mediated transport mechanisms that are regulated by LRRK2.

Recent work indicates that the subcellular co-localization of PPM1H with specific RABs strongly influences levels of RAB phosphorylation^54^. Regulation of RAB-mediated pathways in neurons are therefore determined by the balance of LRRK2 and PPM1H activities at each specific membrane compartment, in ways that are difficult to predict from whole-cell pRAB levels alone. While our results show that both p.R1441H KI and *PPM1H* KO affect SVP transport and synaptic protein distribution, the effect size was generally more pronounced in *PPM1H* KO neurons (Figure 3F,G; Figure S2F). Notably, PPM1H has been shown to strongly localize to the Golgi^13, 54^, suggesting that its loss may cause more striking effects on protein sorting and cargo loading at the TGN. Further work could reveal how relative LRRK2-PPM1H abundance at specific subcellular membrane compartments differentially regulates other RAB-mediated pathways.

While we observed that RAB3A phosphorylation causes strong impairment of the RAB3A-GDI1 interaction (Figure 5A,C), we observed only moderate impairment of RAB3A-MADD (Figure 4E-I) and RAB3A-RAB3GAP (Figure 5A,D) binding, and no impairment of RAB3A-RIM2 (Figure S3A) and RAB3A-synapsin (Figure S3B) interaction. Together, these results predict that RAB3A phosphorylation has a spectrum of effect sizes on effector binding properties, and may alter the probability of successful binding in a manner depending on the specific interfaces of protein-protein interaction. Attempts to model the RAB3A-MADD and RAB3A-GDI1 complexes with AlphaFold-Multimer^40, 41, 55–57^ proved challenging, yielding only low confidence models. However, AlphaFold-Multimer generated a relatively high-confidence model of the complex between RAB3A and RAB3GAP1 (catalytic subunit of RAB3GAP) (Figure S4C; interface pTM + pTM score 0.794)^40, 41, 55–57^. Modeling the RAB3A-MADD complex may be complicated by the presence of intrinsically disordered regions in the large MADD protein (UniProt Q8WXG6). However, previous work showed that the N-terminal 161 residues of MADD are necessary and sufficient for binding to GTP-bound RAB3 isoforms (Figure S4A)^16^. Moreover, MADD’s motor-binding death domain region is found toward its C-terminus^16, 18^. RABs are phosphorylated by LRRK2 at their characteristic switch II domain, which changes conformation in response to nucleotide binding in order to allow for interaction with effectors or regulatory proteins^3, 58^. Thus, phosphorylation of RAB3A likely disrupts interaction between the switch II region of RAB3A and the N-terminus of MADD, without interrupting MADD’s ability to bind kinesin-3 (Figure S4A). Further work is required to elucidate the order of events and kinetics by which pRAB3A interrupts loading of SVP cargo onto the MADD-kinesin motor complex. However, we observed that overexpression of GTP-locked RAB3A rescued anterograde SVP flux in p.R1441H KI iNeurons (Figure 4B-D), suggesting that increasing levels of active RAB3A in this system is sufficient to restore appropriate levels of the RAB3A-MADD-KIF1A/Bβ complex. This indicates that the increased fraction of phosphorylated RAB3A induced by p.R1441H KI reduces the abundance of eligible RAB3A required for the initiation of SVP transport.

Here, we primarily focused on the transport dynamics of the anterograde population of SYP+ vesicles, for which the mechanism of rapid, highly processive transport is known^22^. These results demonstrate that the major effect of RAB3A phosphorylation is on flux of the anterograde SYP+ population. The transport dynamics of the retrograde SYP+ population are not as well-characterized. Across multiple studies in mammalian neurons, we have observed it to be more heterogeneous, and overall less numerous, rapid, and processive than the anterograde population^22, 36^ (Aiken and Holzbaur, 2023; manuscript in preparation). Recent work from our group has identified that SV proteins (including SYP and SYB2) make up a substantial proportion of autophagic cargoes^53^. Thus, a fraction of the retrograde SYP+ population we observe are likely to be synaptic proteins engulfed in AVs and thus moving processively in the retrograde direction along the axon.

Consistent with decreased anterograde SVP flux, our results using p.R1441H KI and *PPM1H* KO iNeurons also show decreased accumulation of SVP-associated proteins in a temporally-controlled model of presynaptic site formation (Figure 3D-G). Given the limited effective lifespan of SVs and the need for continuous replenishment of SVPs at presynapses^21^, it is predicted that RAB hyperphosphorylation and the resulting decrease in anterograde SVP flux would be detrimental to the size and health of the SV pool, especially the readily-releasable and recycling pools. However, the specific ramifications of our findings on synaptic transmission are challenging to disentangle from other roles of LRRK2 activity at the presynaptic site. Despite numerous studies using multiple model systems of LRRK2 hyperactivity or loss (reviewed extensively by Pischedda and Piccoli^59^), no consensus has been reached on the effect of LRRK2 on SV exocytosis. LRRK2 has been reported to interact with or regulate actin, synapsin I, SNAP25, syntaxin, NSF, endophilin A, dynamin, auxillin, and synaptojanin, all of which contribute to the SV cycle^59–68^. Furthermore, LRRK2 substrates RAB3A^49–51, 69, 70^, RAB5^71–73^, and RAB35^74, 75^ have been implicated in membrane trafficking within the presynapse. The relative balance of LRRK2 and PPM1H activity at the presynapse will likely regulate some of these interactions, but others may involve scaffolding domains of LRRK2 that are not believed to be directly kinase-dependent^63, 76^. RAB-mediated pathways at the presynapse may therefore be good candidates to be differentially affected by hyperactive LRRK2. Ultimately, the effect of pathogenic LRRK2 mutations on synaptic transmission is likely an integrated function of these different interactions, with more work needed to uncover how these intersecting pathways may contribute to development of non-motor symptoms in PD.

## ACKNOWLEDGEMENTS

We thank Mariko Tokito for assistance in the cloning of plasmid constructs. We thank Michael Ward (National Institutes of Health), Erika Lara Flores (National Institutes of Health), and Bill Skarnes (The Jackson Laboratory) for expertise in utilizing resources from the iPSC Neurodegeneration Initiative (iNDI). Cartoon schematics were created in part with BioRender.com. Protein structures in Figure 2 were supplied from UniProt.org under the Creative Commons Attribution 4.0 International (CC BY 4.0) License (PDB: 7LHT for LRRK2, 7L4J for PPM1H). AlphaFold and AlphaFold-Multimer (accessed through COSMIC^2^ Gateway) were used for structural models and are cited in the main text. *PPM1H* KO iPSCs used in this study were previously generated by C. Alexander Boecker^12^. Source plasmid DNA for HA-RIM2 was a gift from Ruben Bierings (Erasmus University Medical Center). We thank C. Alexander Boecker, Juliet Goldsmith, and Elizabeth Gallagher for helpful insights and discussions.

This work was supported by the National Institutes of Health (1 F31 NS124249-01 and T32-AG-000255 to D.D., F32 NS117672 to J.A., and R35 GM126950 to E.L.F.H.) and the Michael J. Fox Foundation (MJFF-021130, MJFF-15100, and MJFF-019411 to E.L.F.H.). The E.L.F.H. laboratory is funded by the joint efforts of The Michael J. Fox Foundation for Parkinson’s Research (MJFF) and the Aligning Science Across Parkinson’s (ASAP) initiative. MJFF administers the grant ASAP-000350 on behalf of ASAP and itself.

## DATA AND RESOURCE AVAILABILITY

### Lead Contact

Further information and requests for resources and reagents should be directed to and will be fulfilled by the lead contact, Erika L. F. Holzbaur.

### Materials Availability

Unique reagents generated in this study are available from the Lead Contact with a completed Materials Transfer Agreement. Plasmids generated in this study have been deposited to Addgene (identifier numbers listed in resources table).

### Data Availability

- Primary data that is presented in this study has been deposited in Zenodo and repository and are publicly available as of the date of publication. These can be accessed using the Digital Object Identifier 10.5281/zenodo.8156734.
- Any additional information required to reanalyze the data reported in this paper is available from the lead contact upon request.

## AUTHOR CONTRIBUTIONS

Conceptualization, D.D., J.A., and E.L.F.H.; methodology, D.D. and J.A.; investigation, D.D. and J.A.; writing – original draft, D.D., J.A., and E.L.F.H.; writing – review and editing, D.D., J.A., and E.L.F.H.; funding acquisition, D.D., J.A., and E.L.F.H.; supervision, E.L.F.H.

## METHOD DETAILS

### Plasmids and reagents

Plasmids and reagents used are detailed in Table 1, along with Addgene identification numbers. Antibodies used are detailed in Table 2, along with application and dilution. CMV HA-RIM2 was derived from GST-RIM2-RBD which was a gift from Ruben Bierings (Erasmus University Medical Center). This construct contains the first 411 amino acids of RIM2, which contains the RAB binding domain^77^.

**Table 1.** Plasmids and reagents.

| REAGENT or RESOURCE | SOURCE | IDENTIFIER |
| --- | --- | --- |
| **Recombinant DNA** |  |  |
| Plasmid: PB-TO-hNGN2 | Gift from iPSC Neurodegenerative Disease Initiative (iNDI) & Michael Ward | Addgene plasmid #172115 |
| Plasmid: piggyBac™ transposase vector | Transposagen | N/A |
| Plasmid: PGK mScarlet-Synaptophysin | This paper | Addgene #206145 |
| Plasmid: PGK EGFP-RAB3A-WT | This paper | Addgene #206146 |
| Plasmid: PGK EGFP-RAB3A-Q81L | This paper | Addgene #206147 |
| Plasmid: PGK EGFP-RAB3A-T86E | This paper | Addgene #206148 |
| Plasmid: PGK EGFP-RAB3A-T86A | This paper | Addgene #206149 |
| Plasmid: CMV HA-MADD | This paper; GST-RIM2-RBD was a gift from Ruben Bierings (Erasmus University Medical Center)<sup>77</sup> | Addgene #206150 |
| Plasmid: CMV HA-RIM2 | This paper | Addgene #206151 |
| Plasmid: CMV SNAP-Synapsin | This paper | Addgene #206152 |
| Plasmid: pBI-BFP-NL1 | <sup>36</sup> (Aiken and Holzbaur, 2023; manuscript in preparation) | Addgene #206153 |
| **Chemicals, Peptides, and Recombinant Proteins** |  |  |
| MLi-2 | Tocris | Cat# 5756 |
| DMSO | Sigma-Aldrich | Cat# D2650 |
| Matrigel Growth Factor Reduced | Corning | Cat# 354230 |
| Essential 8 Medium | Thermo Fisher | Cat# A1517001 |
| ReLeSR | Stemcell Technologies | Cat# 05872 |
| Accutase | Sigma-Aldrich | Cat# A6964 |
| ROCK Inhibitor Y-27632 | Selleckchem | Cat# S1049 |
| Knockout Serum Replacement | Thermo Fisher | Cat# 10828010 |
| DMEM/F-12, HEPES | Thermo Fisher | Cat# 11330032 |
| N2 Supplement | Thermo Fisher | Cat# 17502048 |
| Non-essential Amino Acids (NEAA) | Thermo Fisher | Cat# 11140050 |
| GlutaMAX | Thermo Fisher | Cat# 35050061 |
| Doxycycline | Sigma-Aldrich | Cat# D9891 |
| Poly-L-Ornithine | Sigma-Aldrich | Cat# P3655 |
| BrainPhys Neuronal Medium | Stemcell Technologies | Cat# 05790 |
| Laminin | Corning | Cat# 354232 |
| BDNF | PeproTech | Cat# 450-02 |
| NT-3 | PeproTech | Cat# 450-03 |
| B27 Supplement | Thermo Fisher | Cat# 17504-044 |
| Lipofectamine Stem Transfection Reagent | Thermo Fisher | Cat# STEM00003 |
| Microcystin-LR | Sigma-Aldrich | Cat# 475815 |
| Halt Protease and Phosphatase Inhibitor Cocktail | Thermo Fisher | Cat# 78442 |
| 5-Fluoro-2'-deoxyuridine | Sigma-Aldrich | Cat# F0503 |
| Uridine | Sigma-Aldrich | Cat# U3003 |
| DMEM (Dulbecco's Modified Eagle's Medium) | Corning | Cat# MT10-013-CV |
| FuGENE 6 | Promega Corp. | Cat# E2692 |
| GFP-Trap Magnetic Particles M-270 | ChromoTek | Cat# gtd |
| GFP-Trap Magnetic Agarose | ChromoTek | Cat# gtma |
| Lambda ($\lambda$) protein phosphatase | New England BioLabs | Cat# P0753S |
| **Critical Commercial Assays** |  |  |
| BCA Protein Assay Kit | Thermo Fisher | Cat# 23225 |
| Plasmid Maxi Kit | QIAGEN | Cat# 12163 |
| **Experimental Models: Cell Lines** |  |  |
| Human: KOLF2.1J WT iPSCs | B. Skarnes (Jackson Laboratories, Connecticut) | RRID: CVCL_B5P3 |
| Human: KOLF2.1J LRRK2-R1441H iPSCs | B. Skarnes (Jackson Laboratories, Connecticut) | N/A |
| Human: KOLF2.1J *PPM1H* KO iPSCs | This paper | RRID: CVCL_C7TY |
| **Software and Algorithms** |  |  |
| FIJI (Release 2.9.0) | NIH, USA | http://fiji.sc,<br>RRID:SCR_002285 |
| Prism 9 | GraphPad | https://www.graphpad.com/scientific-software/prism/,<br>RRID:SCR_002798 |
| RStudio: Integrated Development for R (2021.09.2 Build 382) | RStudio Team | http://www.rstudio.com/,<br>RRID:SCR_000432 |
| R package: nlme | Pinheiro J, Bates D, R Core Team | http://CRAN.R-project.org/package=nlme,<br>RRID:SCR_015655 |
| Matlab R2022a | MathWorks | https://www.mathworks.com/products/matlab.html,<br>RRID:SCR_001622 |
| Volocity | PerkinElmer | https://www.perkinelmer.com,<br>RRID:SCR_002668 |
| VisiView 5.0.0.24 | Visitron | https://www.visitron.de/products/visiviewr-software.html,<br>RRID:SCR_022546 |
| LI-COR Image Studio | LI-COR | https://www.licor.com/bio/image-studio/,<br>RRID:SCR_015795 |
| Adobe Illustrator 2022 | Adobe | https://www.adobe.com/products/illustrator.html,<br>RRID:SCR_010279 |
| BioRender | BioRender | https://biorender.com/,<br>RRID:SCR_018361 |
| **Other** |  |  |
| 35 mm #1.5 glass bottom imaging dishes | MatTek | Cat# P35G-1.5-20-C |
| ProLong Gold Antifade Mountant | Thermo Fisher | Cat# P36930 |

**Table 2.** Antibodies.

| ANTIBODY | SOURCE | APPLICATION/<br>DILUTION |
| --- | --- | --- |
| **Primary antibodies** |  |  |
| Anti-RAB8A (phospho T72), Rabbit Monoclonal | Abcam (Cat# ab230260, RRID:AB_2814988) | WB @ 1 µg/mL |
| Anti-MAP2, Mouse Monoclonal | Sigma-Aldrich (Cat# MAB3418, RRID:AB_94856) | ICC @ 1:200 |
| Anti-Synaptophysin, Guinea Pig Polyclonal | Synaptic Systems (Cat# 101-004, RRID:AB_1210382) | ICC @ 1:500 |
| Anti-Synaptophysin, Mouse Monoclonal | Santa Cruz (Cat# sc-17750, RRID:AB_628311) | WB @ 1:2000 |
| Anti-Synaptobrevin2 (VAMP2), Rabbit Monoclonal | Cell Signaling Technology (Cat# 13508, RRID:AB_2798240) | ICC @, WB @ 1:250 |
| Anti-RAB3A, Rabbit Polyclonal | Proteintech (Cat# 15029-1-AP, RRID:AB_2177378) | WB @ 1:1000 |
| Anti-Golgin-97, Rabbit Monoclonal | Cell Signaling Technology (Cat# 13192, RRID:AB_2798144) | ICC @ 1:200 |
| Anti-βIII-tubulin | Abcam (Cat# ab7751, RRID:AB_306045) | ICC @ 1:500 |
| Anti-HA, Mouse Monoclonal | BioLegend (Cat# 901501, RRID:AB_2565006) | WB @ 1:2000 |
| Anti-GFP, Chicken Polyclonal | AvesLabs (Cat# GFP-1020, RRID:AB_10000240) | WB @ 1:5000 |
| Anti-SNAP-tag, Rabbit Polyclonal | New England BioLabs (Cat# P9310S, RRID:AB_10631145) | WB @ 1:1000 |
| **Secondary antibodies** |  |  |
| Anti-Rabbit IgG-IRDye 800CW, Donkey Polyclonal | Li-COR Biosciences (Cat# 926-32213, RRID:AB_621848) | WB @ 1:20,000 |
| Anti-Rabbit IgG-IRDye 680RD, Donkey Polyclonal | Li-COR Biosciences (Cat# 926-68073, RRID:AB_10954442) | WB @ 1:20,000 |
| Anti-Mouse IgG-IRDye 800CW, Donkey Polyclonal | Li-COR Biosciences (Cat# 926-32212, RRID:AB_621847) | WB @ 1:20,000 |
| Anti-Chicken IRDye 680RD, Donkey Polyclonal | Li-COR Biosciences (Cat# 926-68075, RRID:AB_10974977) | WB @ 1:20,000 |
| Anti-Rabbit IgG (H+L) Alexa Fluor 555, Goat Polyclonal | Thermo Fisher (Cat# A-32732, RRID:AB_2633281) | ICC @ 1:1000 |
| Anti-Guinea Pig IgG (H+L) Alexa Fluor 488, Goat Polyclonal | Thermo Fisher (Cat# A-11073, RRID: AB_2534117) | ICC @ 1:1000 |
| Anti-Mouse IgG (H+L) Alexa Fluor 647, Goat Polyclonal | Thermo Fisher (Cat# A-32728, RRID:AB_2633277) | ICC @ 1:1000 |

### Piggybac-mediated iPSC-derived neuron differentiation

KOLF2.1J-background WT and *LRRK2*-p.R1441H KI iPSCs were a gift from B. Skarnes (Jackson Laboratories, Connecticut) as part of the iPSC Neurodegenerative Disease Initiative (iNDI) and have been described previously^35^. KOLF2.1J-background *PPM1H* KO iPSCs were generated as described previously^12^. Cytogenetic analysis of G-banded metaphases cells showed a normal male karyotype (Cell Line Genetics). Mycoplasma testing was negative. iPSCs were cultured on plates coated with Growth Factor Reduced Matrigel (Corning) and fed daily with Essential 8 media (Thermo Fisher). To stably express doxycycline-inducible *hNGN2* using a PiggyBac delivery system, iPSCs were transfected with PB-TO-hNGN2 vector (gift from M. Ward, NIH, Maryland) in a 1:2 ratio (transposase:vector) using Lipofectamine Stem (Thermo Fisher). After 72 hours, transfected iPSCs were selected for 48 hours with 0.5 μg/mL puromycin (Takara). Differentiation of iPSCs into iNeurons was performed using an established protocol^35, 38^. In brief, iPSCs were passaged using Accutase (Sigma) and plated on Matrigel-coated dishes in Induction Media (DMEM/F12 supplemented with 1% N2-supplement [GIBCO], 1% NEAA [GIBCO], and 1% GlutaMAX [GIBCO], and containing 2 μg/mL doxycycline. After 72 hours of doxycycline exposure, iNeurons were dissociated with Accutase and cryo-preserved in liquid N_2_. Published protocol can be found on Protocols.io (dx.doi.org/10.17504/protocols.io.e6nvwj54dlmk/v1). Recent work has found that KOLF2.1J iPSCs carry small copy number variants in *ASTN2*^78^. While changes in ASTN2 levels have the potential to alter synaptic strength, this is predicted to occur through postsynaptic rather than presynaptic mechanisms^79^, and is not expected to affect the phenotypes examined here.

### i^3^Neuron differentiation

Pre-i^3^Neuron iPSCs (human iPSCs with an integrated doxycycline-inducible mNGN2 transgene in the AAVS1 safe-harbor locus) were a gift from M. Ward (National Institutes of Health, Maryland) and have been described previously^8, 12, 38, 39^. Cytogenetic analysis of G-banded metaphases cells showed a normal male karyotype (Cell Line Genetics). Mycoplasma testing was negative. Pre-i^3^N iPSCs were cultured on plates coated with Growth Factor Reduced Matrigel (Corning) and fed daily with Essential 8 media (Thermo Fisher). Induction into neuronal fate with doxycycline and cryopreservation of pre-differentiated neurons was performed as described above (“Piggybac-mediated iPSC-derived neuron differentiation”). Published protocol can be found on Protocols.io (dx.doi.org/10.17504/protocols.io.261ge348yl47/v1).

### Culture and transfection of iPSC-derived neurons

Cryo-preserved, pre-differentiated iNeurons (i^3^Neurons or Piggybac-delivered NGN2 neurons) were thawed and plated on live-imaging dishes (MatTek) coated with poly-L-ornithine at a density of 300,000 neurons per dish. For each experimental condition, cells from at least two different batches of induction were used over three or more independent experimental cultures. iPSC-derived neurons were cultured in BrainPhys Neuronal Media (StemCell) supplemented with 2% B-27 (GIBCO), 10 ng/mL BDNF (PeproTech), 10 ng/mL NT-3 (PeproTech), and 1 μg/mL laminin (Corning). 40% of the media was replaced with fresh media twice per week. For Piggybac-delivered NGN2 neurons, 10 μM 5-Fluoro-2′-deoxyuridine and 10 μM uridine were included at the time of plating to prevent survival of mitotic cells. These drugs were removed 24 hours after plating. Live imaging experiments were performed 21 days after thawing pre-differentiated iPSC-derived neurons (DIV21). On DIV18, iPSC-derived neurons were transfected with Lipofectamine Stem (Thermo Fisher) and 1-2.5 μg total plasmid DNA. Immunostaining experiments were performed at DIV14. Published protocol can be found on Protocols.io (dx.doi.org/10.17504/protocols.io.x54v9dj4zg3e/v1).

### Live-cell imaging and motility quantification

iNeurons were imaged on DIV21 in low fluorescence Hibernate A medium (Brain Bits) supplemented with 2% B27, 10 ng/mL BDNF and 10 ng/mL NT-3. Neurons were imaged in an environmental chamber at 37°C. Recordings of mScarlet-SYP+ vesicles for Figures 1 and 4 were acquired on a PerkinElmer UltraView Vox Spinning Disk Confocal system with a Nikon Eclipse Ti inverted microscope, using a Plan Apochromat 60x 1.40 NA oil immersion objective and a Hamamatsu EMCCD C9100-50 camera controlled by Volocity software. Following a scheduled microscope upgrade, live imaging for Figure 2 and Figure 4 was instead performed using a Hamamatsu ORCA-Fusion C14440-20UP camera controlled by VisiView software. Axons were identified based on morphological parameters^39, 80^, and measurements were made to image ∼100-150 µm from the neuronal soma. After identifying this region, one photobleaching cycle was performed with the 405 nm laser for 3 ms/pixel using a ViRTEx Realtime Experiment Control Device. All time lapse recordings were acquired at a frame rate of 5 frames/sec for 5 minutes. Published protocol can be found on Protocols.io (dx.doi.org/10.17504/protocols.io.5jyl8p9mdg2w/v1).

Kymographs of axonal SVPs were generated using the Multiple Kymograph plugin (FIJI). Line width was set to 5 pixels. For flux quantification, anterograde and retrograde vesicle tracks were manually annotated and counted by a blinded investigator. For anterograde velocity quantification, 10-15 representative anterograde vesicle tracks from each kymograph were traced, and velocity was calculated from the average of their slopes.

Figure legends contain the statistical test used and specific p values for each quantification. RStudio version 2021.9.2.382 was used to perform a linear mixed effects model (LME; R package “nlme”). The genotype (or, in MLi-2 experiments, the treatment condition) was treated as the fixed effect. The independent experiment/culture being recorded from was treated as the random effect. For all quantifications, at least three independent experiments were analyzed.

### Immunostaining and quantification

At DIV14, human iNeurons were fixed in 4% paraformaldehyde supplemented with 4% sucrose for 15 minutes, washed four times with PBS, and permeabilized with 0.2% Triton-X in PBS for 15 min. Cells were then blocked for 1 hour with 5% goat serum and 1% BSA in PBS. Neurons were then incubated in primary antibody (see Table 2) diluted in blocking solution at room temperature for 1 hour, washed three times with PBS, and incubated in secondary antibody (see Table 2) diluted in blocking solution for 1 hour at room temperature. After three washes with PBS, coverslips were mounted in ProLong Gold Antifade Mountant (Thermo Fisher). Images were acquired as z stacks at 200 nm step-size using a spinning disk confocal setup as described above. Published protocol can be found on Protocols.io (dx.doi.org/10.17504/protocols.io.8epv5×91ng1b/v1).

For experiments shown in Figure 3A-C, the MAP2 channel was used to select an ROI around the somatic compartment by a blinded investigator. This ROI was used to measure the mean grey value of the SYP and SYB2 signals using sum projections. SYP/SYB2 intensity for each neuron was normalized to the average intensity of the WT neurons from that experimental replicate. Figure legends contain the statistical test used and specific p values for each quantification. RStudio version 2021.9.2.382 was used to perform a linear mixed effects model (LME; R package “nlme”). The genotype was treated as the fixed effect. The independent experiment/culture was treated as the random effect. For all quantifications, at least three independent experiments were analyzed.

For experiments shown in Figure 3D-G, HEK cells transfected 24 hours prior with 6 µL:1 µg mix of FUGENE and pBI-NL1-BFP (bicistronic vector expressing untagged NL1 and cytosolic BFP) were added to DIV13 iNeurons (100K transfected HEK cells added to DIV13 iNeurons cultured in 35 mm imaging dishes). 24 hours after addition of transfected HEK cells, at iNeuron DIV14, cells were fixed for immunocytochemistry in 4% paraformaldehyde supplemented with 4% sucrose and stained as described above (see Table 1 and 2 for antibody information). For analysis, the NL1-BFP channel was used to select an ROI around a NL1+ HEK cell. To determine SYP and SYB2 intensity within presynaptic regions spanning the NL1+ HEK ROI, max-projection images were created and FIJI’s Thresholding tool was used to segment an 8-bit object mask based on the top 1% intensity of the SYP or SYB2 channel. FIJI’s Analyze Particles tool was then used on the object mask redirected to the original SYP or SYB2 image channel to determine intensity values for individual presynaptic puncta. Figure legends contain the statistical test used and specific p values for each quantification. RStudio version 2021.9.2.382 was used to perform a linear mixed effects model (LME; R package “nlme”). The genotype was treated as the fixed effect. The independent experiment/culture and the field of view were treated as the random effects, with field of view nested within experiment. For all quantifications, at least three independent experiments were analyzed.

For experiments shown in Figure S1E-F, the MAP2 channel was used to select an ROI around the somatic compartment by a blinded investigator. This ROI was used to measure the mean grey value of the somatic SYP signal using a sum projection. The “Adjust Threshold” function in ImageJ was used to create a mask on the region of high-intensity golgin-97 signal, and this ROI was used to measure the mean grey value of the SYP signal co-localized with the TGN. Figure legends contain the statistical test used and specific p values for each quantification. RStudio version 2021.9.2.382 was used to perform a linear mixed effects model (LME; R package “nlme”). The genotype was treated as the fixed effect. The independent experiment/culture was treated as the random effects. For all quantifications, at least three independent experiments were analyzed.

### Neuronal lysis and immunoblotting

iNeurons were washed twice with ice cold PBS and lysed with RIPA buffer (50 mM Tris-HCl, 150 mM NaCl, 0.1% Triton X-100, 0.5%deoxycholate, 0.1% SDS, 2x Halt Protease and Phosphatase inhibitor, 2mg/mL microcystin-LR). Samples were centrifuged for 10 min at 17,000 g, and protein concentration of the supernatant was determined by BCA assay. Neuronal proteins (SYP, SYB2, and RAB3A) were resolved on 15% acrylamide gels.

Proteins were transferred to Immobilon-FL PVDF membranes (Millipore) using a wet blot transfer system. Membranes were then stained for total protein using LI-COR Revert 700 Total Protein Stain. Following imaging of total protein stain, membranes were de-stained and blocked for 5 minutes with Bio-Rad EveryBlot blocking buffer. Membranes were incubated with primary antibody diluted in EveryBlot at 4°C overnight. After three washes with TBS (50 mM Tris-HCl [pH 7.4], 274 mM NaCl, 9 mM KCl) with 0.1% Tween-20, membranes were incubated with secondary antibodies diluted in EveryBlot with 0.01% SDS for 1 hr at RT. Following three more washes with TBS with 0.1% Tween-20, membranes were imaged using an Odyssey CLx Infrared Imaging System (LI-COR). Western blots were analyzed with Image Studio Software (LI-COR). When necessary, stripping and reprobing was performed using LICOR NewBlot IR stripping buffer according to manufacturer instructions. Published protocol can be found on Protocols.io (dx.doi.org/10.17504/protocols.io.5jyl8j5zrg2w/v1).

### Cell line culture

HEK293T cells were maintained in DMEM (Corning) with 10% fetal bovine serum (HyClone). Cells were maintained at 37°C in a 5% CO_2_ incubator. Cells were tested for mycoplasma contamination routinely, using MycoAlert detection kit (Lonza, LT07). For co-immunoprecipitation experiments, cells were plated on three 10 cm tissue culture dishes per condition and transfected 24 h before lysis using FuGENE 6 (6-12 μg total plasmid DNA; Promega). Published protocol can be found on Protocols.io (dx.doi.org/10.17504/protocols.io.kxygx3zeog8j/v1).

### Co-immunoprecipitation experiments and quantification

HEK293T cells were lysed 24 h after transfection. For experiments not involving lambda protein phosphatase (λPP), cells were lysed in buffer containing 10 mM Tris-HCl pH 7.5, 150 mM NaCl, and 0.5 mM EDTA, with 0.5% NP-40 and protease inhibitors (1 mM PMSF, 0.01 mg/mL TAME, 0.01 mg/mL leupeptin, 0.001 mg/mL pepstatin A). Lysates were clarified at 10 × g at 4°C for 10 min. GFP-Trap Magnetic Particles M-270 were used for experiments involving MADD due to the protein’s large size; otherwise, GFP-Trap Magnetic Agarose beads were used. 25 μL of bead slurry per experimental condition were washed twice in wash buffer (10 mM Tris-HCl pH 7.5, 150 mM NaCl, 0.5 mM EDTA; with 0.4% Triton-X for MADD pulldowns) for 5 min at 4°C and then equilibrated in lysis buffer for 5 min at 4°C. Beads were incubated with clarified lysate for 1 hour at 4°C under rotating agitation. Following incubation, beads were washed three times for 5 min in wash buffer at 4°C and then resuspended in 60 μL denaturing buffer. The beads were then boiled to release the bound proteins.

For experiments using λPP, cells were lysed in 1x NEBuffer for Protein MetalloPhosphatases (New England BioLabs; supplied with λPP), with 0.5% NP-40 and 0.01 mg/mL leupeptin. Lysates were clarified at 10 × g at 4°C for 10 min. For conditions with λPP, 60 μL of 10 mM MnCl_2_ and 24 μL λPP (2,400 units) were added to clarified lysate for a final reaction volume of 600 μL and incubated at 30°C for 30 minutes. The same was performed for λPP-negative conditions, with 24 μL ddH_2_O instead. After incubation with λPP, incubation with beads was performed as described above. Published protocol can be found on Protocols.io (dx.doi.org/10.17504/protocols.io.6qpvr36o2vmk/v1).

Co-immunoprecipitation was analyzed by Western blot (see “Immunoblotting”). Proteins from the same experiment were processed in parallel and resolved on different acrylamide gels based on protein size: 6% for HA-MADD, 10% for RAB3GAP2, RAB-GDI1, and SNAP-Synapsin, and 12% for EGFP-RAB3A, pT RAB, EGFP vector, and HA-RIM2. Figure legends contain the statistical test used and specific p values for each quantification.

## SUPPLEMENTAL MATERIAL

Supplemental data and legends relating to Figures 1, 3, and 5 can be found in Figures S1-S3. Figure S4 includes AlphaFold and AlphaFold-Multimer predictions relating to Figures 4 and 5.

---

**Figure S1.**
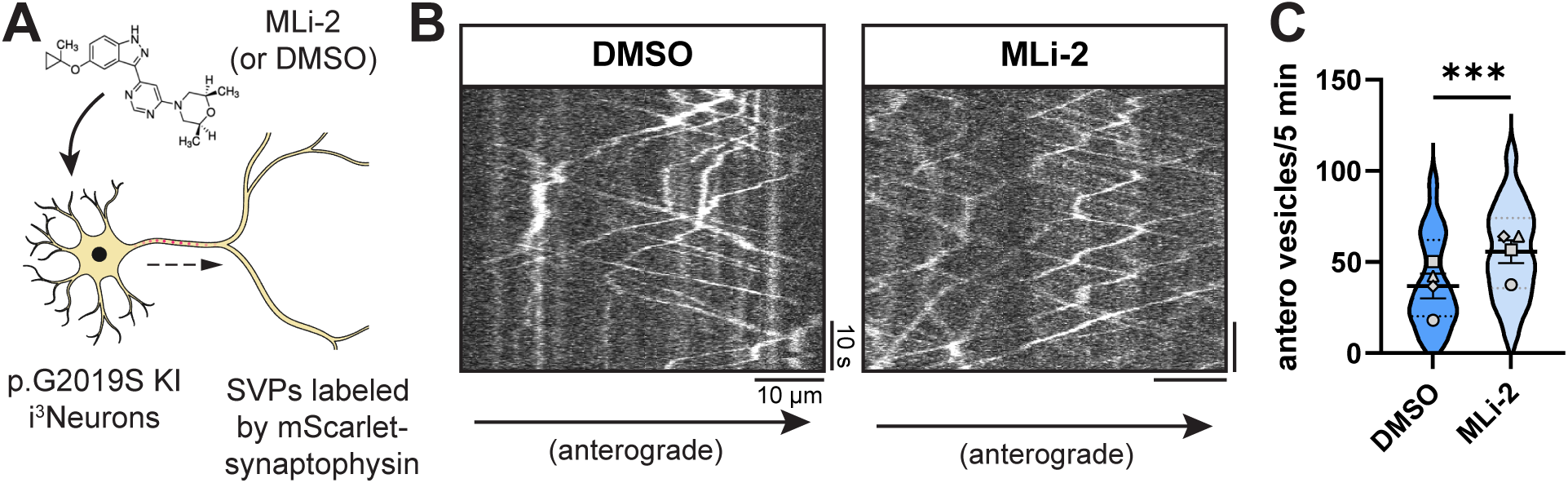
Related to Figure 1. (A) Cartoon depicting p.G2019S KI i^3^Neuron treated overnight with DMSO or 100 nM MLi-2. (B) Kymographs of axonal mScarlet-SYP+ vesicles in p.G2019S KI i^3^Neurons treated with DMSO or MLi-2. (C) Anterograde flux of SYP+ vesicles in p.G2019S KI i^3^Neurons treated with DMSO or MLi-2 (n = 39-41 neurons from 4 independent experiments; ***p<0.001; linear mixed effects model). Scatter plot points indicate the means of four independent experiments, and error bars show mean ± SD of these points.

**Figure S2.**
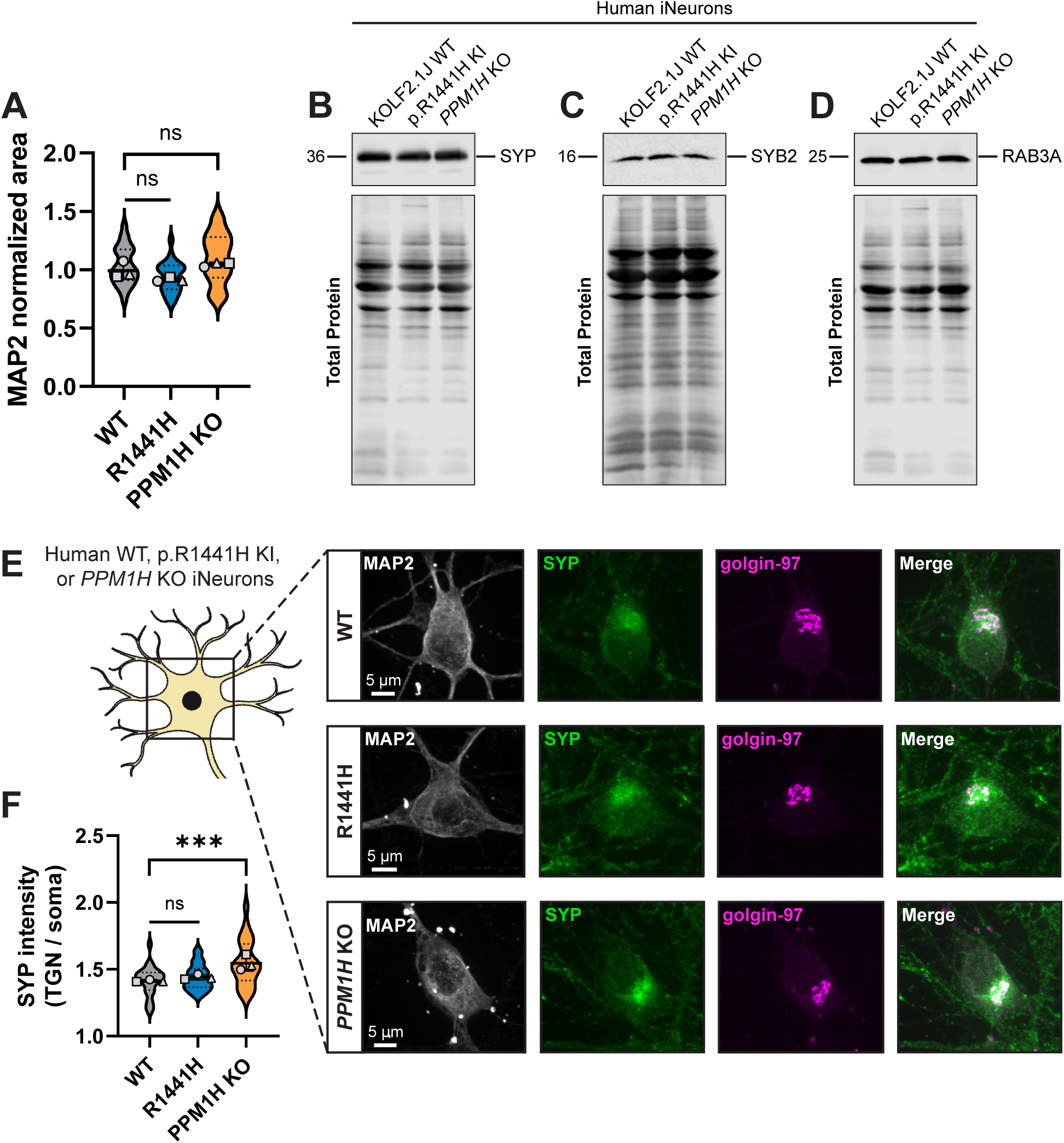
Related to Figure 3. (A) Normalized somal MAP2 area of WT, p.R1441H KI, and *PPM1H* KO iNeurons for dataset shown in Figure 3A-C (n = 24 neurons from 3 independent experiments; ns >0.0540; linear mixed effects model). (B-D) Example total protein stain and immunoblot of SYP (B), SYB2 (C), and RAB3A (D) in DIV21 WT, p.R1441H KI, and *PPM1H* KO iNeurons. (E) Representative images of DIV14 WT, p.R1441H KI, and *PPM1H* KO iNeuron somas, stained for endogenous MAP2, SYP, and golgin-97. (F) Ratio of SYP intensity (mean grey value) co-localized with golgin-97 signal / SYP intensity (mean grey value) of whole soma, in WT, p.R1441H KI, and *PPM1H* KO iNeurons (n = 24 neurons from 3 independent experiments; ns = 0.4638; ***p<0.001; linear mixed effects model). Scatter plot points indicate the means of 3 independent experiments, and error bars show mean ± SD of these points.

**Figure S3.**
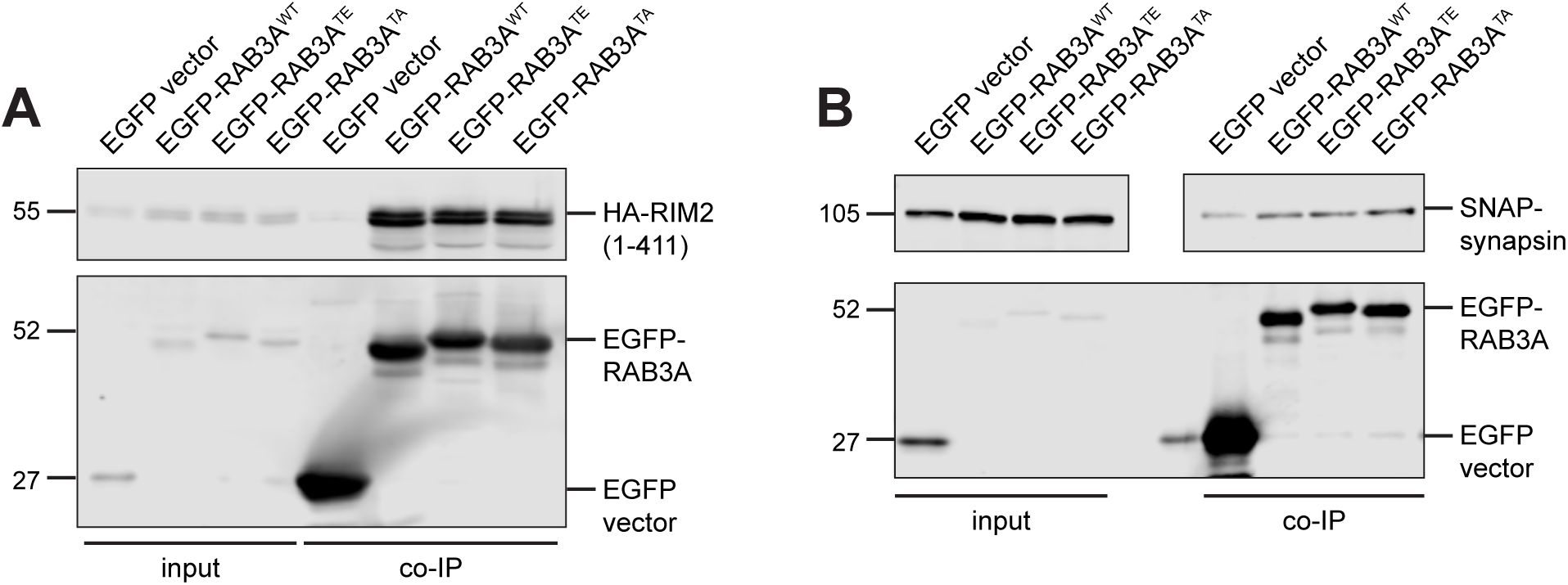
Related to Figure 5. (A) Example immunoblot of RIM2 (first 411 residues) co-immunoprecipitation by RAB3A^WT^, RAB3A^TE^, or RAB3A^TA^, co-expressed in HEK293T cells. (B) Example immunoblot of synapsin co-immunoprecipitation by RAB3A^WT^, RAB3A^TE^, or RAB3A^TA^, co-expressed in HEK293T cells. Upper panel is separated for alignment purposes; no lanes that included sample were excluded. For all co-IP experiments shown, samples were processed and immunoblotted in parallel.

**Figure S4.**
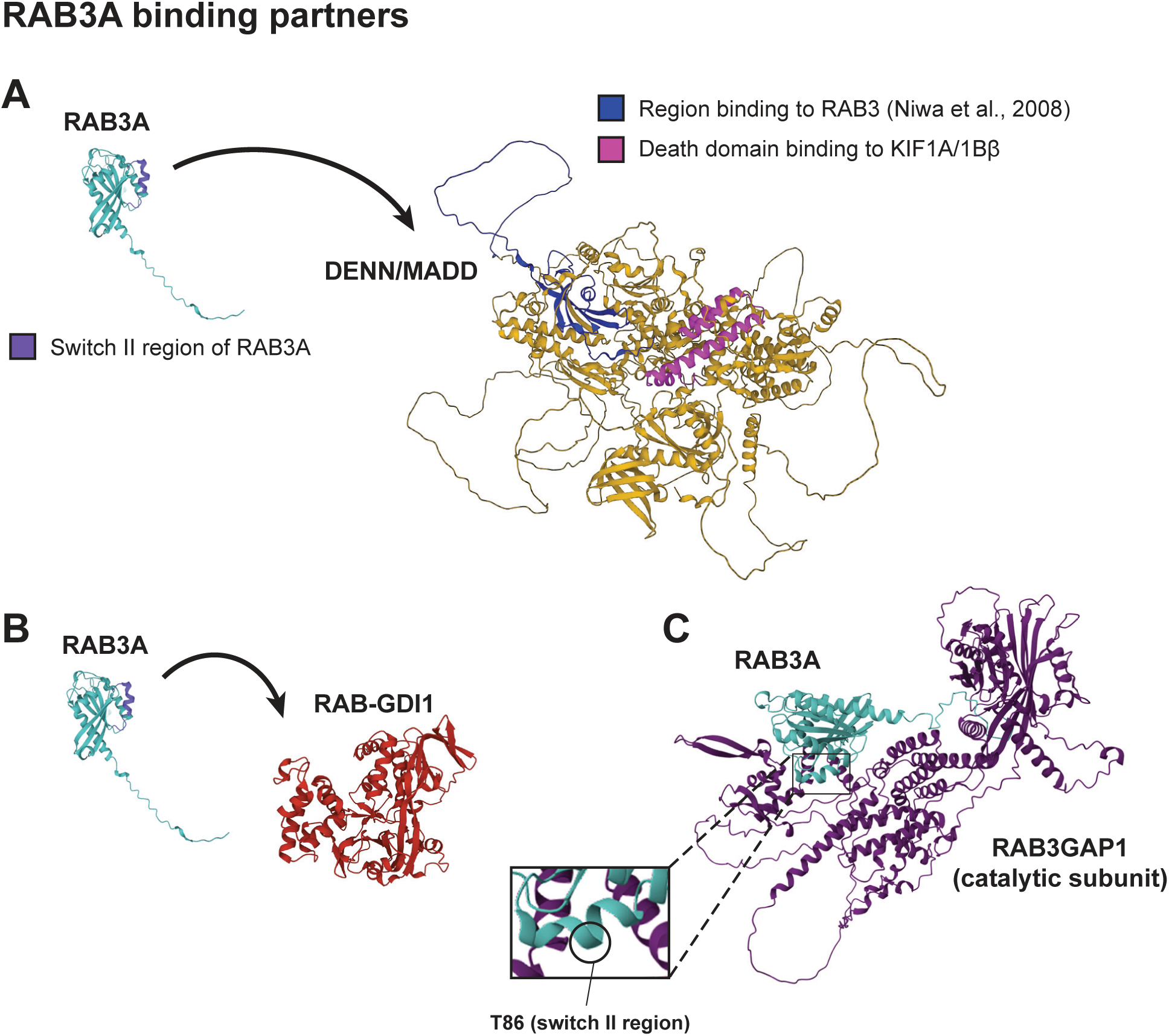
Related to Figures 4 and 5. (A) AlphaFold predictions^40, 41^ of RAB3A and MADD. Left, annotated in purple: putative switch II region of RAB3A^77^. Right, annotated in blue: the N-terminal 161 residues that were previously shown^16^ to be necessary and sufficient for binding to RAB3. Right, annotated in magenta: the death domain toward the C-terminus of MADD that has been shown to be the motor-binding region^16, 18^ (B) AlphaFold predictions^40, 41^ of RAB3A and RAB-GDI1. (C) AlphaFold-Multimer^40, 41, 55–57^ prediction of complex formed by RAB3A and RAB3GAP1, the catalytic subunit of RAB3GAP. ipTM + pTM score for this prediction: 0.79404.

